# Lineage tracing axial progenitors using Nkx1.2CreER^T2^ mice defines their trunk and tail contributions

**DOI:** 10.1101/261883

**Authors:** Aida Rodrigo Albors, Pamela A. Halley, Kate G. Storey

## Abstract

The vertebrate body forms by continuous generation of new tissue from progenitors at the posterior end of the embryo. In mice, these axial progenitors initially reside in the epiblast, from where they form the trunk; and later relocate to the chordo-neural hinge of the tail bud to form the tail. Among them, a small group of bipotent neuromesodermal progenitors (NMPs) are thought to generate the spinal cord and paraxial mesoderm to the end of axis elongation. The study of these progenitors, however, has proven challenging *in vivo* due to their small numbers and dynamic nature, and the lack of a unique molecular marker to identify them. Here, we report the generation of the Nkx1.2CreER^T2^ transgenic mouse line in which the endogenous *Nkx1.2* promoter drives tamoxifen-inducible CreER^T2^ recombinase. We show that Nkx1.2CreER^T2^ targets axial progenitors, including NMPs and early neural and mesodermal progenitors. Using a YFP reporter, we demonstrate that *Nkx1.2*-expressing epiblast cells contribute to all three germ layers, mostly neuroectoderm and mesoderm excluding notochord; and continue contributing neural and paraxial mesoderm tissues from the tail bud. This study identifies the *Nkx1.2*-expressing cell population as the source of most trunk and tail tissues in the mouse; and provides a key tool to genetically label and manipulate this progenitor population *in vivo*.

## Introduction

The vertebrate body forms progressively in a head-to-tail direction from progenitors located at the posterior end of the embryo (reviewed in (Kimelman, 2016; Neijts et al., 2013; Wilson et al., 2009)). In mice, these progenitors initially reside in the epiblast, from where they generate most of the organs of the trunk; and later relocate to the tail bud, from where they generate the tail. The process is thought to be partly fuelled by a small pool of bipotential progenitors with self-renewing capability, the so-called neuromesodermal progenitors (NMPs) (Cambray and Wilson, 2002, 2007; Tsakiridis and Wilson, 2015; Tzouanacou et al., 2009). NMPs give rise to the neural and mesodermal progenitors that form the spinal cord and paraxial mesoderm derivatives (e.g. bones, cartilage, muscle, dermis) of the trunk and tail (reviewed in (Henrique et al., 2015; Steventon and Martinez Arias, 2017)). Lineage tracing studies of small groups of cells in mouse embryos at embryonic day (E) 8.5 have shown that both the neural and paraxial mesoderm tissues of the trunk originate from the epiblast between the node and the anterior primitive streak (the node-streak border or NSB) and the caudal lateral epiblast (CLE) (Cambray and Wilson, 2007; Wymeersch et al., 2016). During the transition from trunk to tail development, this neuro-mesodermal (NM) potential relocates to the chordo-neural hinge (CNH) – the region of the tail bud where the neural tube overlays the posterior end of the notochord (Cambray and Wilson, 2002; McGrew et al., 2008; Wilson and Beddington, 1996). These findings suggest that NMPs first reside within the NSB and CLE and during the transition from trunk to tail development relocate to the CNH.

Molecularly, NMPs have been defined by co-expression of the stem cell and neural transcription factor *Sox2* and the mesodermal transcription factor *T* (*Brachyury*) (Garriock et al., 2015; Tsakiridis et al., 2014; Wymeersch et al., 2016). However, even though cells that co-express *Sox2* and *T* coincide with NM-fated regions, co-expression of *Sox2* and *T* is not a feature unique to NMPs (Wymeersch et al., 2016). Recently, two studies revealed a more complete molecular signature of NMPs and their immediate descendants, early neural and mesodermal progenitors, using single-cell RNA-sequencing technologies (Gouti et al., 2017; Koch et al., 2017). Perhaps not surprisingly, both studies showed that the CLE cell population (Gouti et al., 2017) and cells co-expressing *Sox2* and *T* at E8.5 (Koch et al., 2017) are rather heterogeneous and include, based on their molecular features, NMPs and early neural and mesodermal progenitors. NMPs at E8.5 express *Sox2, T, Nkx1.2, Cdx2* and *Cdx4*, while NMPs at E9.5 and NMPs undergoing lineage choice express NMP marker genes plus *Tbx6* at levels that reflect their fate choice (Gouti et al., 2017; Koch et al., 2017). Accordingly, early mesoderm progenitors express *T* and *Tbx6* and at decreasing levels *Sox2* and *Nkx1.2*, while early neural progenitors express *Sox2*, and at decreasing levels *Nkx1.2* and *T*. Already committed presomitic mesoderm cells express *Msgn1* and *Tbx6* but have repressed *Sox2* and *Nkx1.2*, while neural progenitors express high *Sox2* but have now repressed *Nkx1.2* and mesodermal genes (Gouti et al., 2017; Koch et al., 2017). From these data, it emerges that *Nkx1.2* marks progenitor cells with neural and mesodermal potential. *Nkx1.2* has also been used to identify *in vitro*-derived NMPs (Edri et al., 2018; Gouti et al., 2014; Sasai et al., 2014; Tsakiridis et al., 2014; Verrier et al., 2017). *Nkx1.2*, previously *Sax1* in the chick, is a member of the small NK-l class of homeobox genes. *Nkx1.*2 is widely conserved across species and its expression pattern has been characterised in chick (Rangini et al., 1989; Spann et al., 1994), mouse (Schubert et al., 1995), and zebrafish (Bae et al., 2004). However, the identity of *Nkx1.2*-expressing cells and their contributions to the developing mouse embryo have not been specifically characterised.

Here, we present the first detailed description of the expression pattern of *Nkx1.2* in the mouse embryo and show that it largely overlaps with the posterior growth zone and regions thought to harbour NMPs and early neural and mesodermal progenitors. We describe the generation and characterisation of the Nkx1.2CreER^T2^ transgenic mouse line in which tamoxifen-inducible CreER^T2^ recombinase is driven under the control of the endogenous *Nkx1.2* promoter. We then demonstrate that this line can be used to manipulate gene expression specifically in cells expressing *Nkx1.2* in a temporally-controlled manner. Using a YFP reporter, we trace and define the lineages of the *Nkx1.2*-expressing cell population at different developmental stages and find that this progenitor population is dynamic, changing as development proceeds to supply most tissues of the trunk and tail in the mouse.

## Results

### *Nkx1.2* is expressed in the posterior growth zone throughout body axis elongation

To document in detail *Nkx1.2* expression in the mouse embryo, we carried out whole-mount RNA in situ hybridization and then localised *Nkx1.2* transcripts to specific cell populations in serial transverse sections. As the body develops in a head-to-tail sequence, sections from the posterior end of the embryo represent more undifferentiated structures than more anterior sections. In agreement with a previous report (Schubert et al., 1995), *Nkx1.2* transcripts were first detected around embryonic day (E) 7–7.5 in the NSB as well as in and alongside the primitive streak, in cells of the CLE (Figure 1A-C). This coincides with the emergence of the node and the time and regions in which NMPs first arise during embryonic development (Wymeersch et al., 2016). At E8.5, *Nkx1.2* expression remained highest in epiblast cells in the node region and CLE just posterior to the node (Figure 1D, E, Eb, Ec). *Nkx1.2* was expressed at lower levels in the primitive streak, in cells that ingress to form mesoderm (Figure 1Ec). Anterior to the node, *Nkx1.2* was also expressed in the neural plate, although at lower levels in the midline/floor plate (Figure 1D, E, Ea). The expression pattern and relative levels of *Nkx1.2* in the E8.5 embryo combined with lineage tracing data (Cambray and Wilson, 2007; Wymeersch et al., 2016) support single-cell transcriptomics data suggesting that *Nkx1.2* is highly expressed in NMPs and expressed at lower levels in early neural and mesodermal progenitors (Gouti et al., 2017; Koch et al., 2017). By E9.5 the most anterior *Nkx1.2*-expressing cells have begun to form a neural tube (Figure 1F, Fa, Fb). Posteriorly, transcripts remained in epiblast cells around the closing posterior neuropore but were for the first time detected at lower levels in mesenchymal cells ingressing through the last remnants of the primitive streak as the tail bud forms (Figure 1Fc, Fd). In the tail of E10.5 embryos, *Nkx1.2* transcripts continued to be detected in most newly formed neural tube (Figure 1G, Ga-Gc) and were also found in the CNH region (Figure 1Gb). Here, *Nkx1.2* was expressed in the neural tube and in a mesenchymal cell group continuous with the ventral neural tube, but not in the notochord component of the CNH (Figure 1Gb). Posteriorly, *Nkx1.2* was also expressed in the contiguous dorsal tail bud mesenchyme, albeit at lower levels (Figure 1Gd). Intriguingly, the appearance of this novel mesenchymal *Nkx1.2* domain coincides with the transition from primitive streak to tail bud-driven growth and formation of neural tissue by secondary neurulation, which involves a mesenchymal-to-epithelial transition (Beck, 2015; Lowery and Sive, 2004; Schoenwolf, 1984). At E11.5, *Nkx1.2* transcripts were still detected in the newly formed neural tube and contiguous tail bud mesenchyme (Figure 1H). At all stages, the anterior limit of *Nkx1.2* expression was in the neural tube around the level of the last formed somite (Figure 1D-H). At E12.5, when elongation of the tail is coming to a halt, *Nkx1.2* expression faded away (Figure 1I). Outside of the posterior end of the embryo, *Nkx1.2* transcripts appeared at this stage in a subpopulation of motor neurons in the hindbrain and spinal cord and in the medial longitudinal fascicle of the midbrain (Schubert et al., 1995) (Figure S1).

**Figure 1.**
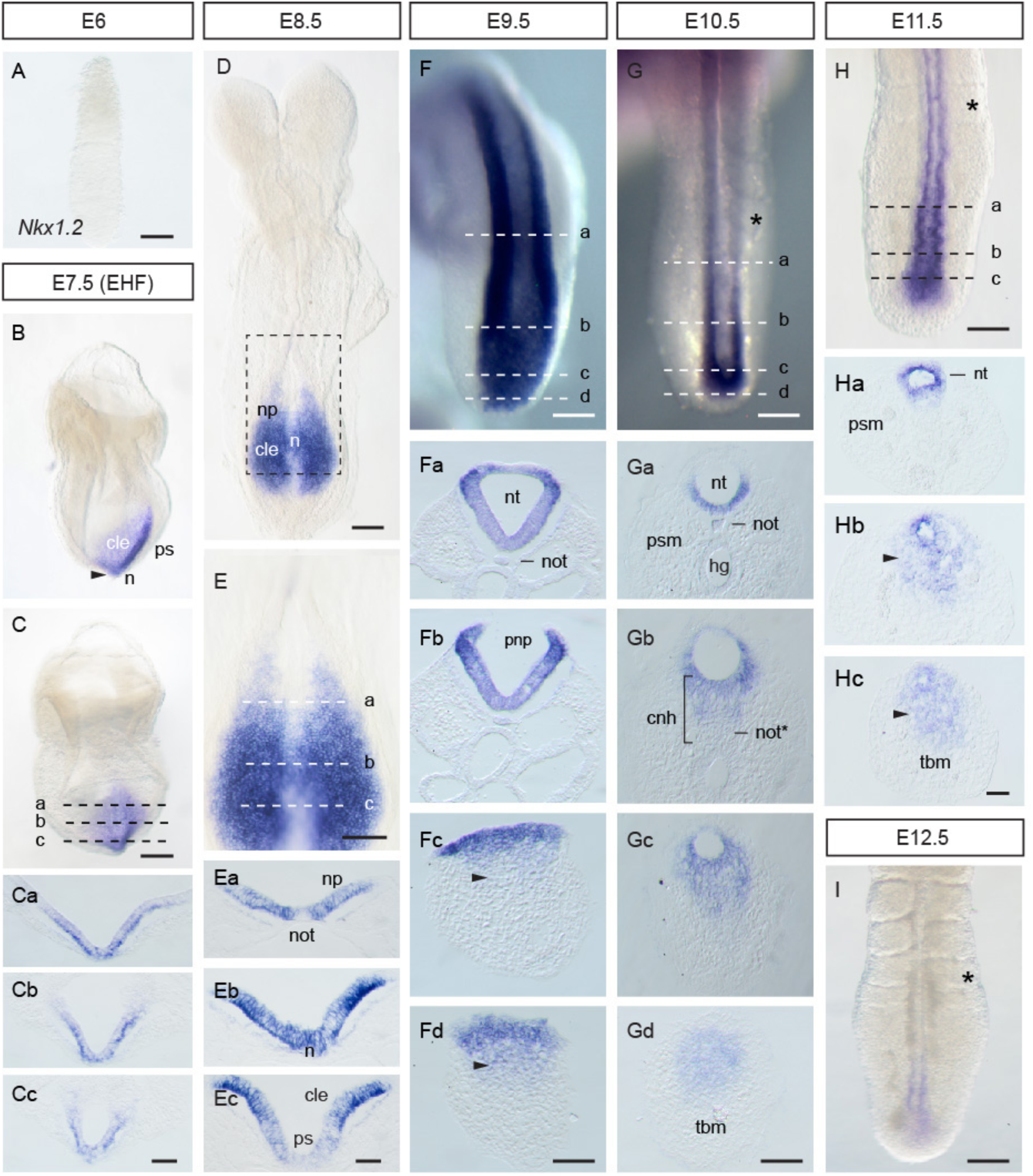
Expression of *Nkx1.2* in the developing mouse embryo. (A) E6.0 embryo (n=4 embryos). (B) Lateral and (C) posterior views of an E7.5 embryo (early head fold) (n=4 embryos). (Ca-Cc) Transverse sections through the regions indicated in C. (D) E8.5 embryo (8–10 somites) and (E) higher magnification of the posterior end of the embryo (n=4 embryos). (Ea-Ec) Transverse sections through the regions indicated in E. (F) Dorsal view of the posterior end of an E9.5 embryo (n=4 embryos). (Fa-Fd) Transverse sections through the regions indicated in F. (G) Dorsal view of the tail end of an E10.5 embryo (n=9 embryos). (Ga-Gd) Transverse sections trough the regions indicated in G. (H) Dorsal view of the tail end of an E11.5 embryo (n=4 embryos). (Ha-Hc) Transverse sections through the regions indicated in H. (I) Dorsal view of the tail end of an E12.5 embryo (n=4 embryos). The arrowheads in (Fc-Fd) and (Hb-Hc) indicate the mesenchymal cell group expressing *Nkx1.2* in the tail bud. The asterisks in (G), (H), and (I) indicate the last formed somite. cle, caudal lateral epiblast; ps, primitive streak; np, neural plate; n, node; np, neural plate; not, notochord; not*, notochord end; nt, neural tube; pnp, posterior neuropore; psm, presomitic mesoderm; hg, hindgut; cnh, chordoneural hinge; tbm, tail bud mesenchyme. Scale bars in whole-mount embryos, 100 µm; scale bars in transverse sections, 50 µm.

Taken together, these data show that *Nkx1.2* expression marks the posterior growth zone and regions thought to harbour NMPs and early neural and mesodermal progenitors throughout body axis elongation.

### *Nkx1.2* regions co-localise with SOX2^+^T^+^ regions fated for neural and mesodermal lineages

NMPs are usually identified *in vivo* by their location and co-expression of the neural transcription factor SOX2 and the mesodermal transcription factor T (Garriock et al., 2015; Tsakiridis et al., 2014; Wymeersch et al., 2016). The relative levels of these two factors correlate with the fate of NMP descendants: neural-fated NMPs gradually increase *Sox2* and decrease *T* expression, while mesoderm-fated NMPs increase *T* and decrease *Sox2* (Gouti et al., 2017; Koch et al., 2017; Wymeersch et al., 2016). To better place NMP cells and their immediate descendants within *Nkx1.2* regions, we carried out SOX2 and T immunofluorescence on transverse sections of mouse embryos. Because embryos display a highly characteristic spatial patterning of tissues along the developing body axis, we used morphological features to align sections with schematic cartoons of the *Nkx1.2* regions defined above (Figure 1). We focused the analysis around the regions known to harbour early and late NMPs – the NSB and CLE at E8.5 and the CNH at E10.5, respectively.

In agreement with previous reports (Garriock et al., 2015; Tsakiridis et al., 2014; Wymeersch et al., 2016), SOX2^+^ T^+^ cells were found at E8.5 at the midline epiblast of the NSB and posteriorly, in the CLE and primitive streak (Figure 2A). Here, T levels were higher in the midline epiblast and primitive streak than in the CLE (Figure 2A) (Wymeersch et al., 2016). Moreover, as recently reported (Javali et al., 2017), TBX6 could be detected in high-T regions: the primitive streak and primitive streak epiblast as well as in the presomitic mesoderm; but not in the NSB epiblast, CLE or neural plate (Figure S2). *Nkx1.2*, however, was expressed across these regions albeit at higher levels in the in the NSB epiblast and CLE than in the primitive streak epiblast (Figure 1Ec and 2A). Taken together, these molecular features suggest that the *Nkx1.2*-expressing cell population at E8.5 includes putative NMPs (SOX2^+^ T^+^ TBX6^-^ *Nkx1.2*^high^) and early neural (SOX^high^ T^-^ TBX6^-^ *Nkx1.2*^low^) and mesodermal (SOX2^low^ T^high^ TBX6^+^ *Nkx1.2*^low^) progenitors.

**Figure 2.**
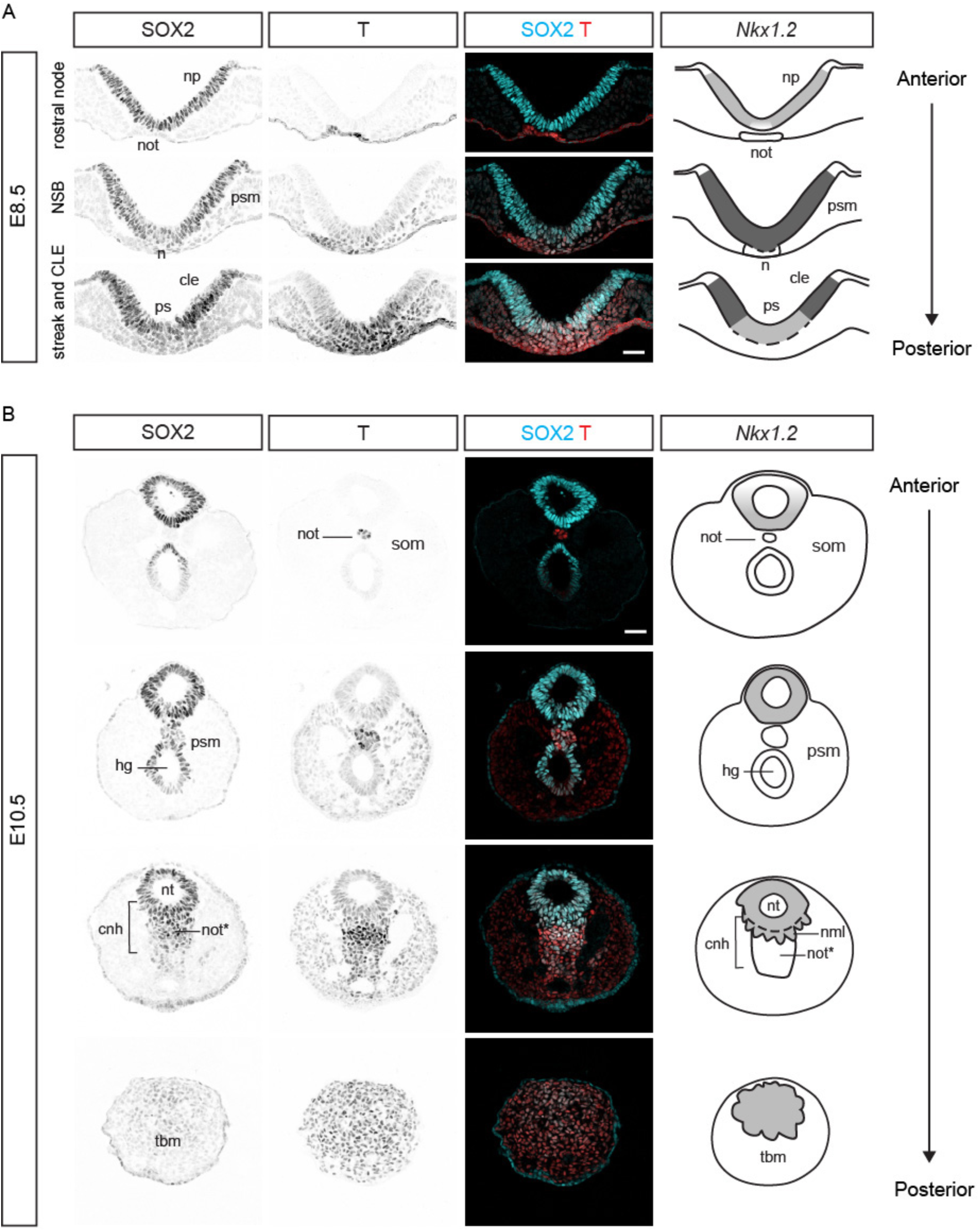
SOX2 and T co-expression within *Nkx1.2* regions. (A) Transverse sections across the rostral node, NSB, and CLE of an E8.5 embryo immunolabelled for SOX2 and T (n=4 embryos). (B) Transverse sections across the tail end of an E10.5 embryo immunolabelled for SOX2 and T (n=7 embryos). The cartoons in (A) and (B) depict the expression pattern of *Nkx1.2* (as shown in Figure 1). The different levels of *Nkx1.2* expression (based on *in situ* hybridisation signal) are represented by different grey intensities (light grey, low; dark grey, high). The dashed lines delineate regions not limited by basement membrane. Abbreviations are the same as in Figure 1. som, somite; nml, neuromesodermal lip. Scale bars, 50 µm.

Between E9.5 and E10.5 NMPs relocate to the CNH region of the tail bud, but their precise location remains unclear. SOX2 and T co-expression is not unique for NMPs in the CNH region: node-derived notochord progenitors and hindgut cells also co-express SOX2 and T, but they are not NMPs (Figure 2B). However, combining SOX2 and T with *Nkx1.2* expression data in the tissue context we could identify as putative NMPs and/or NMP descendants the cells in the dorsal half of the CNH. These included the neural tube and the mesenchymal cells right below the neural tube (Figure 2B). Given the co-expression of neural and mesodermal genes, we propose to name this medial mesenchymal cell population the neuromesodermal lip (NML). In contrast, SOX2^+^T^+^ cells in the ventral half of the CNH express T at higher levels and no or undetectable *Nkx1.2* and thus comprise mostly notochord progenitors (Figure 2B). In agreement with a recent report (Javali et al., 2017), we found low but detectable levels of TBX6 protein in all SOX2^+^ cells in the CNH region, including *Nkx1.2*-expressing cells in the neural tube (higher in the ventral half) and in the NML (Figure S2). This molecular signature of the ventral half of the neural tube and NML (SOX2^+^ T^+^ TBX6^low^ *Nkx1.2*^low^) is consistent with the molecular signature of E9.5 tail bud NMPs proposed by single-cell transcriptomics data (Gouti et al., 2017) and what Koch and colleagues proposed to be NMPs undergoing lineage choice (Koch et al., 2017). Posterior to the CNH, cells of the tail bud mesenchyme also co-express SOX2 and T proteins and low levels of *Nkx1.2* transcripts, resembling cells in the primitive streak epiblast at E8.5 (Figure 2). Lineage tracing of dorsal tail bud mesenchyme (Cambray and Wilson, 2002; McGrew et al., 2008) and the molecular signature of this cell population, including TBX6 (Figure S2), suggest that *Nkx1.2*-expressing cells in the tail bud mesenchyme (*Nkx1.2*^low^ SOX2^low^ T^high^ TBX6^+^) are early mesoderm progenitors. As expected, presomitic mesoderm cells express high levels of T and TBX6 and neither SOX2 or *Nkx1.2* (Figure 2B, Figure S2) (Chalamalasetty et al., 2014; Gouti et al., 2017).

Taken all together, these data provide a refined map of the *in vivo* location of NMPs and NMP immediate descendants throughout body axis elongation, and put forward *Nkx1.2* as reliable marker for these dynamic progenitor populations.

### Generation of the Nkx1.2CreER^T2^ mouse line

To label and manipulate specifically *Nkx1.2*-expressing cells and thus potentially NMPs and early neural and mesodermal progenitors in a temporally-controlled manner, we set out to generate a transgenic mouse line in which the expression of CreER^T2^ recombinase is driven under the control of the *Nkx1.2* promoter. The CreER^T2^ sequence was knocked in to the *Nkx1.2* locus in C57BL/6 ES cells using standard targeting methods by Taconic Biosciences. The strategy used to generate the conditional transgenic mouse is summarised in Figure 3. Taconic provided two breeding pairs of heterozygous C57BL/6-Nkx1.2^tm2296(Cre-ER(T2))Arte^ (Nkx1.2CreER^T2^) mice. These mice carried a puromycin-expressing cassette flanked by FLP sites, which was removed upon crossing to Flp-expressing mice. The resulting animals were then bred to homozygosity to establish a breeding colony. Knocking out *Nkx1.2* did not generate a phenotype in either heterozygous or homozygous mice, likely due to functional redundancy with the paralogous gene *Nkx1.1* (Bober et al., 1994) (Frank Schubert and Peter Gruss personal communication). Here, we substantiate this unpublished finding with the maintenance of a homozygous Nkx1.2CreER^T2^ transgenic line for at least 9 generations without deleterious effects.

**Figure 3.**
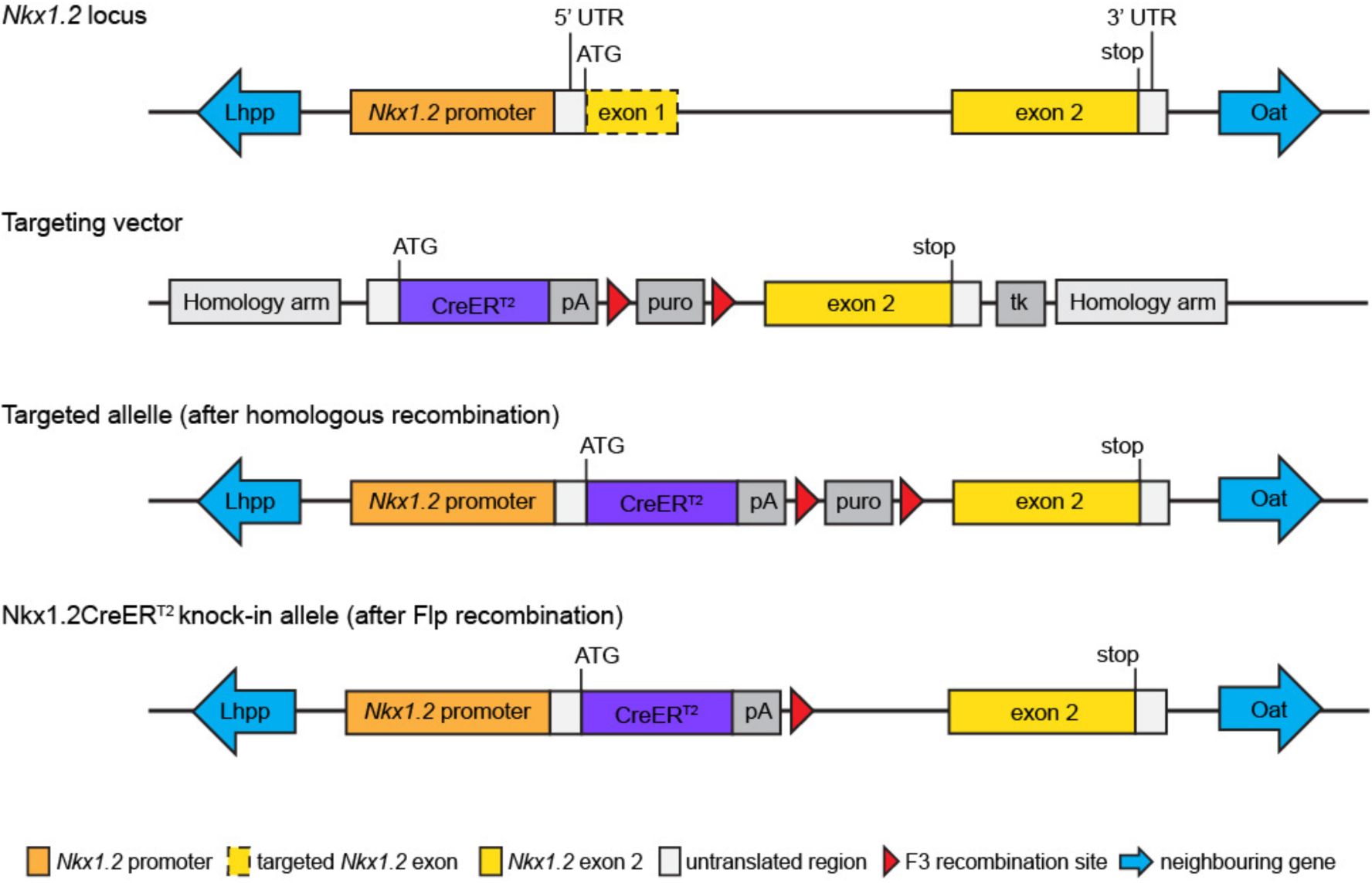
Strategy to knock-in the CreER^T2^ cassette into the *Nkx1.2* locus. *Nkx1.2* locus and targeting vector designed to replace the *Nkx1.2* coding sequence in exon 1 as well as the splice donor site at the junction between exon 1 and intron 1 with a cassette containing the open reading frame (ORF) of CreER^T2^. A polyadenylation site (pA) was inserted 3’ of the CreER^T2^ ORF to terminate transcription. Positive clones were isolated using positive (puromycin, puro) as well as negative (thymidine kinase, tk) selection. Recombinant clones were injected into mouse blastocysts and transferred to mice. The resulting chimeric mice were bred to Flp deleter mice that ubiquitously express Flp recombinase to remove the puromycin selection marker to generate the Nkx1.2CreER^T2^ line.

Next, to fluorescently label *Nkx1.2*-expressing cells in developing embryos, homozygous Nkx1.2CreER^T2^ females were mated with heterozygous or homozygous males harbouring a *loxP*-flanked stop sequence upstream of a EYFP reporter gene under the control of the ubiquitous *ROSA26* promoter (R26R*-*EYFP mice) (Srinivas et al., 2001). In the resulting Nkx1.2CreER^T2^ floxed EYFP mice (Nkx1.2CreER^T2^/YFP), tamoxifen administration leads to CreER^T2^-mediated recombination of the *loxP*-flanked stop sequence and expression of the YFP reporter in *Nkx1.2*-expressing cells and their progeny. We confirmed that Nkx1.2CreER^T2^ faithfully drives transgene expression in the endogenous *Nkx1.2* regions and that only low levels of spontaneous recombination occur in the absence of tamoxifen, but always within canonical *Nkx1.2* regions (Figure S3). Thus, overall, tamoxifen-induced CreER^T2^-mediated recombination leads to faithful YFP-labelling of *Nkx1.2*-expressing cells in the Nkx1.2CreER^T2^/YFP reporter mouse.

### Nkx1.2CreER^T2^/YFP labels SOX2 and T co-expressing cells and their progeny

To establish which cells are labelled with the Nkx1.2CreER^T2^/YFP reporter and whether they include NMPs, we set out to identify YFP^+^ cells based on their location and expression of SOX2 and T. Timed-pregnant Nkx1.2CreER^T2^/YFP mice received tamoxifen either at E7.5 (the onset of *Nkx1.2* expression) or E9.5 (during relocation of axial progenitors to the CNH) to label *Nkx1.2*-expressing cells around these stages and 24 hours later we analysed the posterior growth zone in the epiblast and tail bud. In embryos exposed to tamoxifen at E7.5 and analysed at E8.5, most SOX2^+^ T^+^ cells in the epiblast layer of the NSB and CLE were also YFP^+^. This suggests that Nkx1.2CreER^T2^/YFP labels putative NMPs (Figure 4A). As would then be expected, YFP^+^ cells were also found in the neural plate (SOX2^+^ T^-^), ingressing mesoderm, and paraxial mesoderm (SOX2^-^ T^+^) (Figure 4A). Additionally, a few YFP^+^ cells were found in intermediate and lateral plate mesoderm as well as prospective surface ectoderm (Figure 4A). These findings indicated that the *Nkx1.2*-expressing cell population in the epiblast around E7.5 is heterogeneous, composed of NMPs and early neural and paraxial mesoderm progenitors, lateral plate and intermediate mesoderm progenitors, as well as a few endoderm-fated progenitors. In embryos exposed to tamoxifen at E9.5 and analysed at E10.5, a subset of YFP^+^ cells co-expressed SOX2 and T in the dorsal half of the CNH, including the neural tube and the NML (Figure 4B). YFP^+^ cells were, however, absent from the notochord component of the CNH (SOX2^+^ T^high^ cells) and from hindgut precursors and the hindgut proper (also SOX2^+^T^+^). Besides the CNH, YFP^+^ SOX2^+^ T^+^ cells populated the contiguous tail bud mesenchyme (Figure 4B) and YFP^+^ cells contributed to NMP lineages: most newly formed neural tube and paraxial mesoderm (Figure 5). Overall, using the Nkx1.2CreER^T2^/YFP reporter it is possible to specifically label axial progenitors, including NMPs and their most immediate descendants, at specific developmental stages. This short tracing experiments also suggested that the *Nkx1.2*-expressing cell population is dynamic, changing lineage contributions as development proceeds.

**Figure 4.**
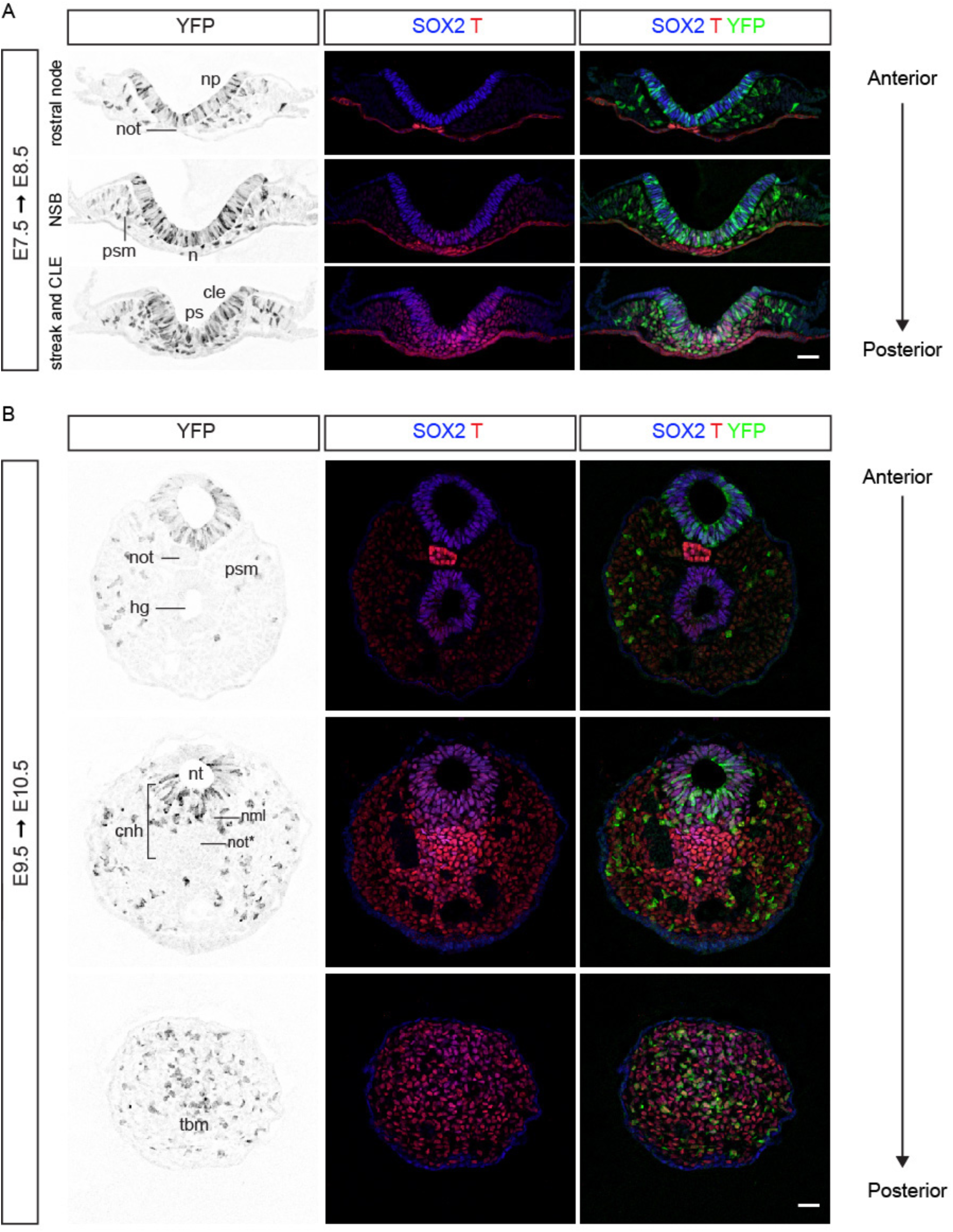
A subset of *Nkx1.2*-expressing cells and/or their progeny express SOX2 and T. (A) Transverse sections through the rostral node, NSB, and CLE of an E8.5 Nkx1.2CreER^T2^ embryo that was exposed to tamoxifen at E7.5 and immunolabelled for SOX2, T, and YFP. (n=7 embryos). (B) Transverse sections through the tail end of an E10.5 Nkx1.2CreER^T2^ embryo that was exposed to tamoxifen at E9.5 and immunolabelled for SOX2, T, and YFP (n=9 embryos). Abbreviations are the same as in Figure 1. nml, neuromesodermal lip. Scale bars, 50 µm.

**Figure 5.**
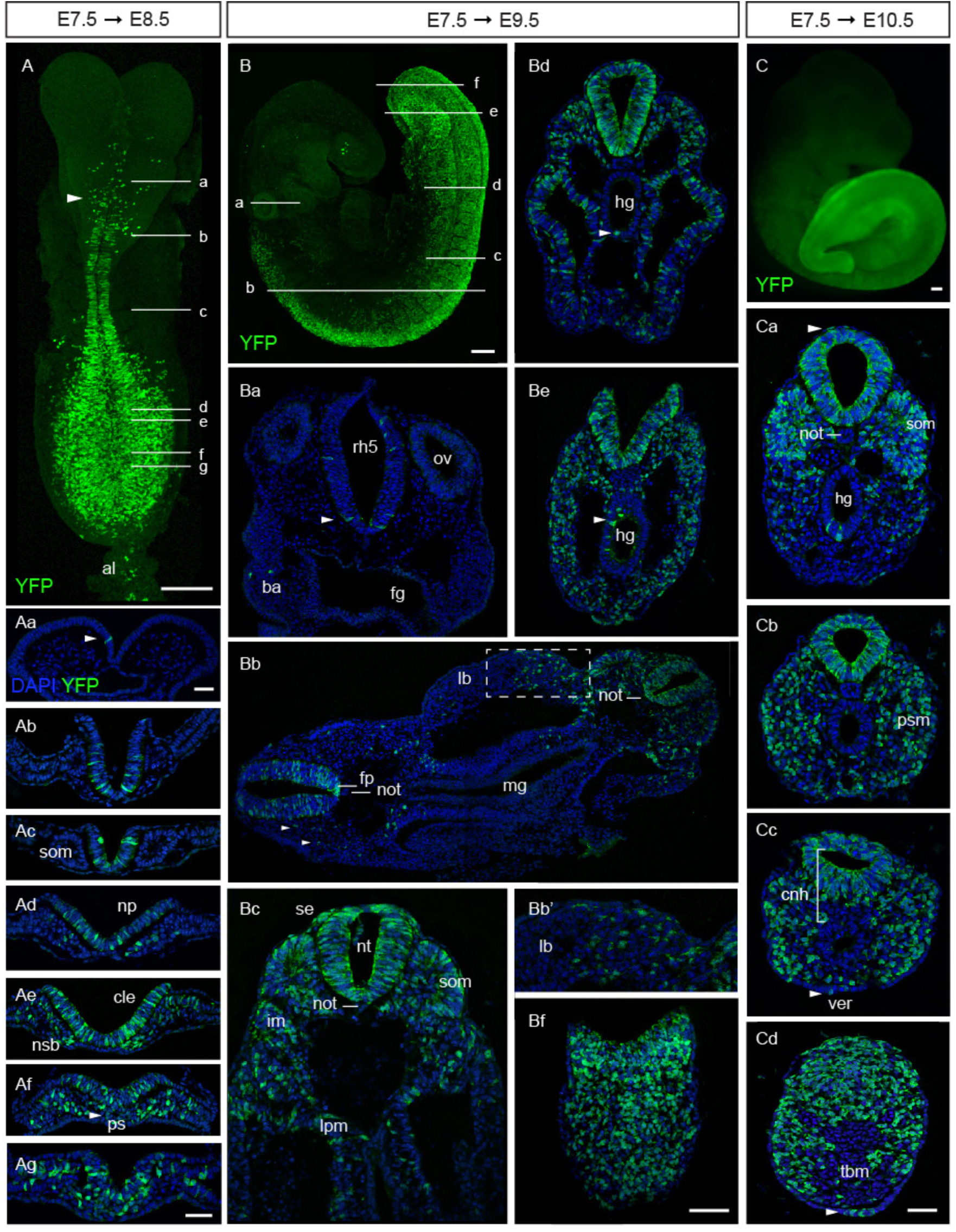
Lineage tracing of cells expressing *Nkx1.2* at E7.5. Timed-pregnant Nkx1.2CreER^T2^ mice received tamoxifen at E7.5 and the contribution of YFP^+^ cells to developing embryos assessed at E8.5, E9.5, and E10.5. Maximum intensity projection (MIP) of an E8.5 embryo immunolabelled for YFP on whole-mount (n=7 embryos). The arrowhead marks the presumptive midbrain/anterior hindbrain boundary. (Aa-Ag) Transverse sections through the regions indicated in A (n=8 embryos). (B) MIP of a E9.5 embryo immunolabelled for YFP on whole-mount (n=12 embryos). (Ba-Bf) Transverse sections through the regions indicated in B (n=8 embryos). (Ba) At the level of the otic vesicles (ov) and rhombomere 5 (rh5) scattered YFP^+^ cells were found in the neural tube (arrowhead). A few YFP^+^ cells populated the branchial arches (ba). The foregut (fg) was, however, always unlabelled. (Bb) Section comprising the anterior (left) and posterior (right) levels of the trunk. YFP^+^ cells located mostly in the floor plate (fp) of the neural tube at more anterior levels, but spanned the dorsoventral extent of the more posterior neural tube. YFP^+^ cells generated neural crest cells (arrowheads), contributed to posterior somites, and to limb bud (lb) mesenchyme. YFP^+^ cells were absent from notochord (not) and midgut (mg). (Bb’) Higher magnification of limb bud mesenchyme in Bb (dashed box). (Bc) YFP^+^ cells contributed to the neural tube (nt), somites (som), intermediate mesoderm (im), lateral plate mesoderm (lpm), and surface ectoderm (se). (Bd, Be) YFP^+^ cells were found frequently in the hindgut (hg) (arrowhead). (Bf) YFP^+^ cells extended to the caudal epiblast and underlying mesenchyme. (C) Widefield fluorescence image of an E10.5 embryo that received tamoxifen at E7.5. Most of the posterior body derived from YFP^+^ cells at this stage (n=7 embryos). (Ca) YFP^+^ cells made most of the neural tube and somites (som) and also contributed to hindgut (hg) endoderm and surface ectoderm (arrowhead), but were absent from the notochord (not). (Cb) Posterior to Ca, most presomitic mesoderm (psm) is YFP^+^. (Cc) Besides in the neural tube and paraxial mesoderm, YFP^+^ cells were found in the NML in the CNH (cnh) region. YFP^+^ cells were also present in the VER (ver, arrowhead). (Cd) The tail bud mesenchyme (tbm) except the ventral-medial cell group was YFP^+^. The images are representative images of each stage and anterioposterior level. Scale bars, 100 µm on whole-mount; 50 µm on transverse sections.

### Early *Nkx1.2*-expressing cells contribute to all three germ layers

To investigate the long-term contribution of early *Nkx1.2-*expressing cells and their progeny to the developing embryo, timed-pregnant Nkx1.2CreER^T2^/YFP mice received a single dose of tamoxifen at E7.5 and the contribution of YFP^+^ cells was assessed in embryos at progressively later developmental stages. Assessment of embryos at E8.5 revealed scattered single cells across the presumptive midbrain/anterior hindbrain as the anterior limit of cells derived from *Nkx1.2*-expressing cells (Figure 5A, Aa). YFP^+^ cells were found contiguously in the neural tube from the presumptive posterior hindbrain (Figure 5A, Ab) and then, more posteriorly, throughout the CLE and primitive streak (Figure 5A, Ad-Ag). YFP^+^ cells were also present in derivatives of the primitive streak: the recently ingressed mesoderm and most recently formed presomitic mesoderm, but were absent from the first 4–5 somites (Figure 5Ad-Ag). At the posterior end of the embryo, a few YFP^+^ cells also contributed to intermediate mesoderm and lateral plate mesoderm compartments as well as the allantois (Figure 5A, 5Af-Ag). YFP^+^ cells were consistently absent from the notochord (Figure 5A, Ac).

A day later, in E9.5 embryos, again a few scattered cells were located in the midbrain and the roof of the anterior hindbrain as well as the developing eye (Figure 5B). The anterior limit of contiguous YFP labelling was now clearly located in the hindbrain just anterior to the otic vesicle in rhombomere 5 (Figure 5B). More posteriorly, YFP^+^ cells were concentrated ventrally, in the floor plate of the spinal cord (Figure 5Ba, Bc). This finding suggests that the floor plate of the trunk spinal cord originates from cells expressing *Nkx1.2*, probably the dorsal layer of the node (see Figure 1Eb), as indirectly suggested by the combined results of earlier cell labelling studies (Beddington, 1994; Sulik et al., 1994). From forelimb levels to the posterior end of the embryo, YFP^+^ cells were found throughout the dorsoventral extent of the neural tube, in somites and their derivatives (Figure 5Bb, Bc). YFP^+^ cells also contributed extensively to intermediate and lateral plate mesoderm (Figure 5B, Bc). Mesenchymal cells derived from the lateral plate mesoderm could be seen migrating in to the limb bud (Figure 5Bc). From forelimb levels, YFP^+^ cells also appeared in the surface ectoderm (Figure 5Bc-Bf) and as streams of neural crest cells emerging from the dorsal neural tube (Figure 5Bc). YFP^+^ cells were absent from the first 4–5 somites, but contributed to both medial and lateral compartments of the posterior-most 11 to 12 somites (Figure 5B, Bb, Bd). YFP^+^ cells did not contribute to the notochord (Figure 5Ba-c), although a few isolated YFP^+^ cells were found in the notochord of one embryo. This finding argues against a common source of floor plate and notochord after E7.5 and agrees with grafting and cell labelling experiments that indicated that the ventral node, which does not express *Nkx1.2* (see Figure 1Eb), is the source of trunk notochord (Beddington, 1994; Brennan et al., 2002; Yamanaka et al., 2007). In all E9.5 embryos examined, YFP^+^ cells were absent from the fore-and midgut (Figure 5Ba-Bc), but frequently found in the hindgut (Figure 5Bd, Be). At the posterior end of the embryo, YFP^+^ cells made most of the posterior neuropore and underlying mesenchyme (Figure 5Bc-Bf).

Overall, these lineage tracing studies show that the majority of descendants of E7.5 *Nkx1.2*-expressing cells contribute to the neural and mesodermal tissues of the trunk, including paraxial, intermediate, and lateral plate mesoderm as well as to the extraembryonic allantois; and that at least some E7.5 *Nkx1.2*-expressing cells and/or their progeny are retained at the growing end of the embryo. Interestingly, a recent study derived NMP-like cells in a dish that resemble the caudal epiblast of the embryo at the time of emergence of the node (around E7.5, coinciding with the emergence of the *Nkx1.2*-expressing cell population) and found that such *in vitro* NMP-like cells possess the potential to differentiate not only into neural and paraxial mesoderm cells but also into intermediate and lateral plate mesoderm (Edri et al., 2018). These findings confirm that such a multipotent progenitor cell population exists in the embryo within the E7.5 epiblast and can be identified by the expression of *Nkx1.2*. In addition to neural and mesodermal tissues, we found descendants of early *Nkx1.2*-expressing cells contributing to neural crest, surface ectoderm, and hindgut endoderm.

Between E9.5 and E10.5, axial progenitors complete trunk formation and begin forming a tail from the tail bud region. In E10.5 Nkx1.2CreER^T2^/YFP embryos exposed to tamoxifen at E7.5, YFP^+^ cells made up most of the neural tube and paraxial mesoderm/somites of the tail (Figure 5C, 5Ca-Cd). Posteriorly, YFP^+^ cells populated the dorsal half of the CNH (Figure 5Cc) and the presomitic mesoderm as well as the tail bud mesenchyme contiguous with these regions (Figure 5Cd). YFP^+^ cells were, however, virtually absent among the cells contiguous with the posterior end of the notochord and the hindgut, in the ventral compartment of the CNH and the ventral tail bud mesenchyme (Figure 5Cd). YFP^+^ cells were found very rarely in the tail notochord and occasionally in the hindgut (Figure 5C, Ca-c). A few YFP^+^ cells contributed to surface ectoderm and to the ventral ectodermal ridge (VER) (Figure 5Cc). This long-term lineage tracing experiment indicates that cells that expressed *Nkx1.2* at E7.5 and/or their progeny persist in the posterior end of the embryo from where they continue generating the neural and paraxial mesoderm tissues of the tail.

### Late *Nkx1.2*-expressing cells continue to make neural and mesodermal tissues from the tail bud

To test whether the *Nkx1.2*-expressing cell population in the tail bud retains neural and paraxial mesoderm potential, timed-pregnant Nkx1.2CreER^T2^/YFP mice received tamoxifen at E10.5 and embryos were assessed 24 or 48hours later. In E11.5 embryos, YFP^+^ cells were indeed found in the neural tube and in the paraxial mesoderm as well as in the tail bud mesenchyme (Figure 6). YFP^+^ cells in the neural tube extended from the tail bud to axial levels right below the hindlimb (opposite to somite ~36), where the transition from trunk to tail development and from primary to secondary neurulation takes place (Shum et al., 2010). YFP^+^ cells in the paraxial mesoderm were found more posteriorly, in newly generated presomitic mesoderm (Figure 6). This different distribution of YFP^+^ cells in the neural tube and paraxial mesoderm likely reflects the broader expression of *Nkx1.2* in the neural tube (Figure 1G). *Nkx1.2*-expressing cells labelled at E9.5 did not contribute to surface ectoderm or the hindgut (Figure 4B) and, as expected, cells labelled at E10.5 did not contribute to these tissues in E11.5 embryos either (Figure 6). The latter is consistent with lineage tracing of single cells (Lawson et al., 1991; Lawson and Pedersen, 1992) and of cell groups (Tam and Beddington, 1987; Wilson and Beddington, 1996) in the early primitive streak which all indicate that the gut endoderm lineage derives from epiblast cells prior to E8.5. Analysis of E12.5 embryos 48hours after tamoxifen administration confirmed that cells expressing *Nkx1.2* at E10.5 contribute to both the neural tube and paraxial mesoderm/somites of the tail, while being retained at the posterior end of the embryo (Figure 6B). These findings indicate that the *Nkx1.2*-expressing cell population retains the potential to generate neural and paraxial mesoderm tissues and to self-renew until the end of body axis elongation. An outstanding question, however, is whether the *Nkx1.2-*expressing cell population undergoes progressive lineage restrictions or whether different axial progenitors are “recruited” in-demand as the embryo elongates.

**Figure 6.**
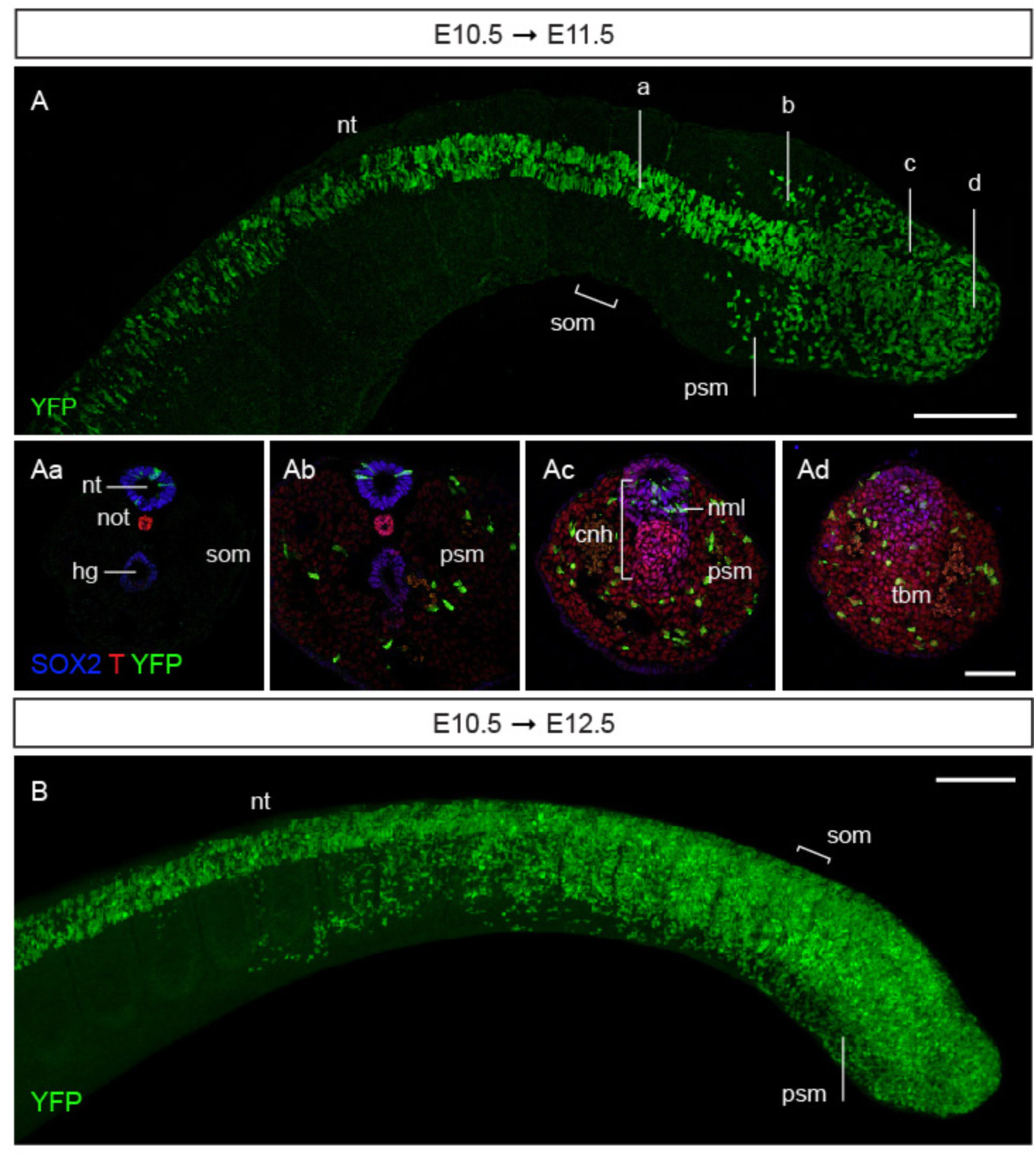
Lineage tracing of cells expressing *Nkx1.2* at E10.5. Timed-pregnant Nkx1.2CreER^T2^ mice received tamoxifen at E10.5 and the contribution of YFP^+^ cells to developing embryos assessed at E11.5 (A) and E12.5 (B). (A) Dorsal view (maximum intensity projection, MIP) of the tail of a E11.5 embryo immunolabelled for YFP on whole-mount (n=7 embryos). (Aa-Ad) Transverse sections representative of the levels indicated in (A) and immunolabelled for SOX2, T, and YFP (n=9 embryos). YFP^+^ cells contributed to the secondary neural tube (see Aa-Ad), presomitic mesoderm (Ab-Ad), and tail bud mesenchyme (Ad). (B) Side view (MIP) of the tail of a E12.5 embryo immunolabelled for YFP on whole-mount (n=9 embryos). Abbreviations are the same as in Figure 1. Scale bars, 100 µm on whole-mount; 50 µm on transverse sections.

Taken all together, this study shows that *Nkx1.2*-expressing cells comprise a heterogeneous and changing cell population with self-renewing capability that generates most trunk and tail tissues in the mouse.

## Discussion

Axial progenitors, including NMPs, have proven challenging to study in the developing mouse embryo because they comprise a relatively small (Wymeersch et al., 2016), dynamic (this study and (Aires et al., 2016; Gouti et al., 2017; Javali et al., 2017; Koch et al., 2017)), and transient cell population. Lineage tracing studies using dye labelling or grafting small groups of cells can be imprecise because of the close proximity between various types of progenitors and extensive cell mixing in the rapidly growing embryo. A complementary approach is to use transgenic reporter mice to label and manipulate specifically progenitors *in vivo*. An important limitation of this, however, is that the promoters/Cre driver lines used so far have not been shown to specifically target NMPs or just axial progenitors (Garriock et al., 2015; Javali et al., 2017; Jurberg et al., 2013; Wymeersch et al., 2016).

### *Nkx1.2* marks axial progenitors including NMPs and early neural and mesodermal progenitors

This study puts forward *Nkx1.2* as a consistent marker of the posterior growth zone in the elongating mouse embryo – where axial progenitors, including the NMPs and early neural and mesodermal progenitors reside.

We have shown that *Nkx1.2* expression overlaps with NMP regions: with the NSB epiblast and adjacent CLE during trunk development and later with the CNH of the tail bud (Cambray and Wilson, 2007; Wymeersch et al., 2016). *Nkx1.2* is also expressed at lower levels around these regions, in early neural and mesodermal progenitors (Figure 1). Our characterisation of the *Nkx1.2* region in the E8.5 embryo has identified cell populations with molecular signatures of early NMPs and early neural and mesodermal progenitors (Chalamalasetty et al., 2014; Gouti et al., 2017; Koch et al., 2017). Interestingly, *Nkx1.2* expression at the posterior end of the E8.5 embryo much resembles the region of transcriptional activity of the *Sox2 N1* enhancer, which is active in NMPs (Takemoto et al., 2011) and it seems, progenitor regions with NM potential (Wymeersch et al., 2016). The *Sox2 N1* enhancer and *Nkx1.2* expression are both promoted by Fgf signalling (Delfino-Machín et al., 2005; Sasai et al., 2014; Takemoto et al., 2011), which acts together with Wnt signalling to regulate and maintain the NMP pool (Garriock et al., 2015; Jurberg et al., 2014; Wymeersch et al., 2016). This further suggests that the expression of *Nkx1.2* marks the axial progenitor state. Importantly, once the tail bud has formed cells that co-express *Nkx1.2*, SOX2 and T now all express TBX6, and so have the molecular signature that distinguishes primitive streak epiblast at E8.5 (Javali et al., 2017) (see Figure S2). This has recently been suggested to represent a transition state in NMPs undergoing lineage choice (Gouti et al., 2017; Javali et al., 2017; Koch et al., 2017) and would be in line with the loss of the pluripotency factor *Oct4* in progenitors relocated to the tail bud (Aires et al., 2016). Because TBX6 represses the activity of the *Sox2 N1* enhancer associated with the NMP state, it would be interesting to see whether this regulatory element is repressed in the tail bud or if it remains active in the cells that express *Nkx1.2*.

Together these findings indicate that the composition of the *Nkx1.2*-expressing cell population changes as axial progenitor populations emerge and most distinctly, when part of the *Nkx1.2* expressing cell population is internalised in the tailbud.

### *Nkx1.2*-expressing cells generate most of the tissues of the trunk and tail

To access genetically the *Nkx1.2-*expressing cell population in the embryo and at different developmental stages, we have generated the Nkx1.2CreER^T2^ transgenic mouse line. By crossing this Nkx1.2CreER^T2^ mouse to a R26R-EYFP reporter (Srinivas et al., 2001) we have traced the contributions of the *Nkx1.2*-expressing cell population to trunk and tail development. Our long-term lineage tracing experiments showed that the *Nkx1.2*-expressing cell population in the E7.5 epiblast includes progenitors for most trunk tissues: neuroectoderm and all mesoderm tissues (paraxial, intermediate and lateral plate mesoderm) except axial mesoderm (notochord); and some progenitors for the posterior-most gut endoderm (hindgut) (Figure 5). In these experiments, cells that had expressed *Nkx1.2* remained at the posterior end of the embryo and relocated to the tail bud, confirming their self-renewing capability and thus the presence of long-term axial progenitors within the *Nkx1.2*-expressing cell population. Lineage tracing *Nkx1.2*-expressing cells in the tail bud showed that this progenitor pool continues to generate the neural and paraxial mesoderm tissues of the tail; but not notochord, hindgut, or surface ectoderm (Figure 6). This is in agreement with earlier lineage tracing studies labelling or grafting small groups of cells in this region (Cambray and Wilson, 2007; McGrew et al., 2008). The lineage restriction of the *Nkx1.2*-expressing cell population likely reflects the response of progenitors to the changing environment and tissue requirements in the elongating embryo. For example, when the transition to tail development is taking place, lateral and intermediate mesoderm progenitors differentiate/are used up to generate the hindlimbs and ventral lateral mesoderm of the trunk (Jurberg et al., 2013).

Taken together, these findings indicate that *Nkx1.2*-expressing cells comprise a self-renewing and changing progenitor cell population that generates most tissues of the trunk and tail. The Nkx1.2CreER^T2^ transgenic mouse line provides now the opportunity to dissect how these progenitors form the posterior body in the developing mouse embryo. For example, crossing the Nkx1.2CreER^T2^ mouse line with a conditional multicolour reporter such as the R26R-Confetti mouse (Snippert et al., 2010) may prove a useful tool to accurately investigate the contributions of single axial progenitors. Crossing the Nkx1.2CreER^T2^ mouse line, with conditional knock-in or knock-out mice is already providing new mechanistic insights into how this important cell population is directed to form trunk and tail tissues at the right place and time (Mastromina et al., 2017; Nikolopoulou et al., 2017; Rolo et al., 2016).

## Materials and methods

### Mice and tamoxifen administration

Wild-type CD-11 and C567BL/6 J mouse strains and transgenic lines Nkx1.2CreER^T2^, R26R-EYFP (Srinivas et al., 2001), Nkx1.2CreER^T2^/YFP were maintained on a 14-hours light/10-hours dark cycle. For timed matings, the morning of the plug was considered E0.5. To make a tamoxifen stock solution, tamoxifen powder (Sigma T5648) was dissolved in vegetable oil to a final concentration of 40 mg/ml and sonicated to bring to solution. The tamoxifen stock solution was stored at −20 ºC for up to three months. At various stages of pregnancy, Nkx1.2CreER^T2^/YFP females were given a single 200 µl dose of tamoxifen (of the 40 mg/ml stock) via oral gavage. Mice were monitored for 6 hours and when required sacrificed following schedule 1 of the Animals (Scientific Procedures) Act of 1986. Embryos were dissected in ice-cold PBS and fixed in ice-cold 4% paraformaldehyde (PFA) for 2 hours (for immunofluorescence) or overnight (for RNA *in situ* hybridization). Gastrula embryos were staged according to (Downs and Davies, 1993) and at later stages by standard morphological criteria. All animal procedures were performed in accordance with UK and EU legislation and guidance on animal use in bioscience research. The work was carried out under the UK project license 60/4454 and was subjected to local ethical review.

### Genotyping

Genotyping by standard methods was performed to maintain the homozygous line using the following PCR conditions: 95°C, 5 min and then 95°C, 30s; 60°C, 30 s; 72°C, 1 min for 35 cycles followed by 72°C, 10 min. A DNA quality control and a test reaction were carried out in parallel for the KI allele, the wild-type (WT) allele, and the Flpe deleter (TG) using the following primer pairs:

KI primer 1: 5’ACGTCCAGACACAGCATAGG 3’, primer 2: 5’TCACTGAGCAGGTGTTCAGG 3’ (fragment size 279 bp); QC primer 3: 5’GAGACTCTGGCTACTCATCC 3’; primer 4: 5’CCTTCAGCAAGAGCTGGGGAC 3’ (fragment size 585 bp); WT primer 5:

5’CAAGGTTTATTGGTAGCCTGG 3’, primer 6: 5’TGAGCCAGTCAGAGTTGTGG 3’ (fragment size 176 bp); QC primer 7: 5’GTGGCACGGAACTTCTAGTC, primer 8: 5’CTTGTCAAGTAGCAGGAAGA 3’ (fragment size 335 bp); TG primer 9: 5’GGCAGAAGCACGCTTATCG 3’, primer 10: 5’GACAAGCGTTAGTAGGCACAT 3’ (fragment size 343 bp); QC primer 3 as above, QC P4 as above (fragment size 585 bp).

### RNA *in situ* hybridization

Standard methods were used to carry out mRNA in situ hybridisation in wild type CD-1 and C57BL/6 J (Charles River) mouse embryos (Wilkinson and Nieto, 1993). The *Nkx1.2* plasmid was kindly provided by Frank Schubert. This probe includes the homeobox domain and the 3’ half of the gene (nucleotides 504–1057).

### Immunofluorescence and imaging

Embryos were permeabilised by dehydration in an increasing methanol series (25% methanol/PBS, 50% methanol/PBS, 75% methanol/PBS, 100% methanol), then stored in 100% methanol at −20 ºC; or bleached in 3% H_2_O_2_/methanol and gradually rehydrated in PBS in preparation for immunofluorescence. For whole-mount immunofluorescence, whole embryos were blocked in PBS/0.1% Triton X-100 (PBST) and 10% normal donkey serum (NDS) for 4 hours and incubated with primary antibodies in PBST/NDS (1:500) overnight at 4 ºC. After incubation with primary antibodies, embryos were washed extensively in PBST (throughout the day or to the next day) and then incubated with secondary antibodies (1:500) and and DAPI (1 mg/ml stock solution diluted 1:500) in PBST/10% NDS overnight at 4 ºC. Embryos were then washed extensively for 24 hours and prepared for clearing. For BABB (2:1 benzyl alcohol:benzyl benzoate) clearing, embryos were first dehydrated in an increasing methanol series (25% methanol/PBS, 50% methanol/PBS, 75% methanol/PBS, 100% methanol. 5 minutes each), then put in 1:1 (v/v) methanol:BABB for 5 minutes and twice in BABB for clearing. BABB-cleared embryos were mounted in BABB for imaging. For immunofluorescence on cryosections, embryos were cryoprotected in 30% sucrose/PBS overnight at 4 ºC, mounted in agar blocks (1.5% agar/5% sucrose/PBS), and frozen on dry ice. 16 µm-thick sections were cut on a Leica CM1900 cryostat, mounted on adhesion slides, and dried for several hours at room temperature. Slides were then washed three times in PBST and blocked in PBST/10% NDS at room temperature. After at least 1 hour, sections were incubated with primary antibodies in PBST/10% NDS overnight at 4 ºC. After several PBST washes, sections were incubated with secondary antibodies and DAPI in PBST/10% NDS for 2 hours at room temperature or overnight at 4 ºC. After several PBST washes, slides were mounted with SlowFade Gold antifade mountant (Invitrogen, S36936) for imaging.

Primary antibodies and working dilutions used were: chicken anti-GFP (Abcam, ab13970; 1:500), goat anti-GFP (Abcam, ab6673; 1:500), rabbit anti-SOX2 (Millipore, AB5603; 1:500), goat anti-SOX2 (Immune Systems, GT15098; 1:500), goat anti-Brachyury/T (R&D Systems, AF2085; 1:500), goat anti-TBX6 (R&D Systems, AF4744; 1:200). Secondary antibodies used, all at a1:500 working dilution, were: donkey anti-chicken Alexa Fluor 488 (Abcam, ab150173), donkey anti-goat Alexa Fluor 488 (Life Technologies, A11055), donkey anti-rabbit Alexa Fluor 568 (Life Technologies, A10042), donkey anti-goat Alexa Fluor 647 (Life Technologies, A21477).

Whole-mount embryos and tissue sections were imaged on a confocal laser scanning microscope Leica TCS SP8 in the Dundee Imaging Facility. Tissue sections were in some cases scored on a Leica DB fluorescence microscope or with a DeltaVision imaging system.

### Methodology

The sample size of each experiment is reported in the respective figure legend. In all cases, *n* reflects the number of embryos analysed per experiment. All experiments were repeated at least twice (so embryos are from at least two independent litters). No statistical methods were used to predetermine sample size. The experiments were not randomised and the investigators were not blinded during to the group allocation or outcome assessment.

## Acknowledgements

We thank the University of Dundee WBRUTG and the Dundee Imaging Facility for technical assistance and advice, Val Wilson and Moisés Mallo for prompt and insightful comments on the preprint version of this manuscript, and to Alwyn Dady and Ioannis Kasioulis for comments on this final version. Creation of the Nkx1.2CreER^T2^ line and initial characterisation was supported by MRC grant G1100552 (P.A.H. and K.G.S.). This work was also supported by a Wellcome Trust Senior Investigator Award WT102817 to K.G.S. Microscopes used for imaging were purchased with support from a Multi-User Equipment grant WT101468 from the Wellcome Trust.

## Author contributions

Conceptualization: A.R.A. and K.G.S; Methodology: A.R.A., P.A. H., K.G.S; Validation: A.R.A. and P.A.H. Formal analysis: A.R.A., P. A. H., K.G.S.; Investigation: A.R.A., P.A. H., K.G.S; Resources: K.G.S; Data Curation: A.R.A.; Writing – original draft preparation: A.R.A., K.G.S; Writing – review and editing: A.R.A., P.A. H., K.G.S; Visualization: A.R.A; Supervision: K.G.S; Project administration: K.G.S; Funding acquisition: K.G.S.

The authors declare no competing interests.

